# A CRISPRi screen of essential genes reveals that proteasome regulation dictates acetic acid tolerance in *Saccharomyces cerevisiae*

**DOI:** 10.1101/2021.04.06.438748

**Authors:** Vaskar Mukherjee, Ulrika Lind, Robert P. St. Onge, Anders Blomberg, Yvonne Nygård

## Abstract

CRISPR interference (CRISPRi) is a powerful tool to study cellular physiology under different growth conditions and this technology provides a means for screening changed expression of essential genes. In this study, a *Saccharomyces cerevisiae* CRISPRi library was screened for growth in medium supplemented with acetic acid. Acetic acid is a growth inhibitor challenging the use of yeast for industrial conversion of lignocellulosic biomasses. Tolerance towards acetic acid that is released during biomass hydrolysis is crucial for cell factories to be used in biorefineries.

The CRISPRi library screened consists of >9,000 strains, where >98% of all essential and respiratory growth-essential genes were targeted with multiple gRNAs. The screen was performed using the high-throughput, high-resolution Scan-o-matic platform, where each strain is analyzed separately. Our study identified that CRISPRi targeting of genes involved in vesicle formation or organelle transport processes led to severe growth inhibition during acetic acid stress, emphasizing the importance of these intracellular membrane structures in maintaining cell vitality. In contrast, strains in which genes encoding subunits of the 19S regulatory particle of the 26S proteasome were downregulated had increased tolerance to acetic acid, which we hypothesize is due to ATP-salvage through an increased abundance of the 20S core particle that performs ATP-independent protein degradation. This is the first study where a high-resolution CRISPRi library screening paves the way to understand and bioengineer the robustness of yeast against acetic acid stress.

**IMPORTANCE:** Acetic acid is inhibitory to the growth of the yeast *Saccharomyces cerevisiae*, causing ATP starvation and oxidative stress, which leads to sub-optimal production of fuels and chemicals from lignocellulosic biomass. In this study, where each strain of a CRISPRi library was characterized individually, many essential and respiratory growth essential genes that regulate tolerance to acetic acid were identified, providing new understanding on the stress response of yeast and new targets for the bioengineering of industrial yeast. Our findings on the fine-tuning of the expression of proteasomal genes leading to increased tolerance to acetic acid suggests that this could be a novel strategy for increasing stress tolerance, leading to improved strains for production of biobased chemicals.

## INTRODUCTION

Systematic profiling of relationships between genotypes and phenotypes provides novel understanding of fundamental biology and suggests leads for improving strains for various biotechnology applications. Quantitative phenotyping of different collections of strains with systematic genetic perturbations, such as the yeast deletion collection (1), the yeast GFP clone collection (2) or yeast overexpression collections (3, 4) has allowed construction of yeast regulatory network models. Nonetheless, the function of a large number of genes remains unknown and many known genes may have more functions yet to be discovered. Notably, even small perturbations in expression of genes can lead to large phenotypic changes (5).

In recent years, the CRISPR interference (CRISPRi) technology has been demonstrated as a very efficient tool to alter gene regulation (6). This technology exploits an RNA-guided, endonuclease-dead Cas9 (dCas9), or other CRISPR-associated proteins, for controlled downregulation of genes by directing dCas9-fusions to their promoter region (7). This allows alteration of expression of essential genes, as partial loss-of-function phenotypes can be induced by conditional expression of *dCas9* and the target-gene specific guide RNA (gRNA). Furthermore, as the strength of expression alteration is greatly dependent on the efficiency and positioning of the gRNA, one can study a gradient of repression by testing multiple gRNA sequences for each target gene (8, 9). Based on this technology, several CRISPRi strain libraries were constructed for many species, including *Saccharomyces cerevisiae* (9-13).

In the first CRISPRi library constructed for yeast (12), transcriptional interference was achieved with an integrated dCas9-Mxi1 repressor (14) and a tetracycline-regulatable repressor (TetR) that controls the expression of the gRNA (8). In this strain collection of roughly 9,000 strains, nearly 99% of the essential and 98% of the respiratory growth essential genes have been targeted with up to 17 gRNAs per target gene (12). Recently, the construction and phenotypic screening of CRISPR technology-based *S. cerevisiae* libraries have been demonstrated to be very efficient to identify bioengineering genetic candidates to increase production of β-carotene or endoglucanase (15), regulate polyketide synthesis (16) or improve tolerance to furfural (11) or lignocellulose hydrolysate (13).

Lignocellulose hydrolysates contain not only fermentable sugars but also various amounts of other compounds, including furfural, different weak acids, and phenolic compounds, that inhibit yeast growth (reviewed by Jönsson et al. (17)). Among these compounds, toxicity by acetic acid is one of the most limiting factors for the production of alternative fuels and chemicals from lignocellulosic biomass using *S. cerevisiae*. Acetic acid is formed during hydrolysis of biomass and is inhibitory to yeast even at low concentrations (17). Tolerance to acetic acid is a very complex trait, where many genetic elements play together to control the phenotype (reviewed by Fernández-Niño et al. (18)). As a result, rational designing of acetic acid tolerant strains is particularly challenging (19).

In this study, a CRISPRi library (12) was used to screen essential and respiratory growth essential genes for roles in providing tolerance towards acetic acid in *S. cerevisiae*. The library was characterized using the automated high-resolution and high-throughput Scan-o-matic platform (20), where each strain is analyzed separately for its growth rate on solid medium. A set of strains with interesting acetic acid growth profiles were verified in liquid medium and the repression of some of these genes was verified by qPCR. The library enabled us to confirm previously known genes involved in the response to acetic acid and to identify several novel genes the regulation of which could be altered to increase tolerance towards acetic acid and thereby improve production hosts for production of biocommodities from lignocellulosic biomass.

## RESULTS

### High-throughput phenomics of the CRISPRi strains

To identify genes involved in tolerance of *S. cerevisiae* to acetic acid, we performed a high-throughput growth screen of a CRISPRi library (9,078 strains) targeting essential and respiratory growth essential genes (12) using the Scan-o-matic system (20) (Fig. 1). The screens were independently duplicated, in total resulting in >27,000 images, and the image analysis generated >42 million data-points and >140,000 growth curves. Our large-scale screen showed rather good repeatability (Fig. 2A). Linear regression, taking all strains into account, showed that 22% (co-efficient of determination i.e. R^2^ = 0.22, F-test P-value < 2.2e-16) of the phenotypic variability between the two independent screens could be explained by the linear model (Pearson correlation coefficient *r* = 0.47). However, taking only the strains with distinct phenotypes into account i.e., statistically significant acetic acid sensitive or tolerant strains (Fig. 2A), 79% (R^2^ = 0.79, F-test P-value < 2.2e-16) of the phenotypic variability between the two independent experiments could be explained by the linear model (*r* = 0.89).

**Fig. 1.**
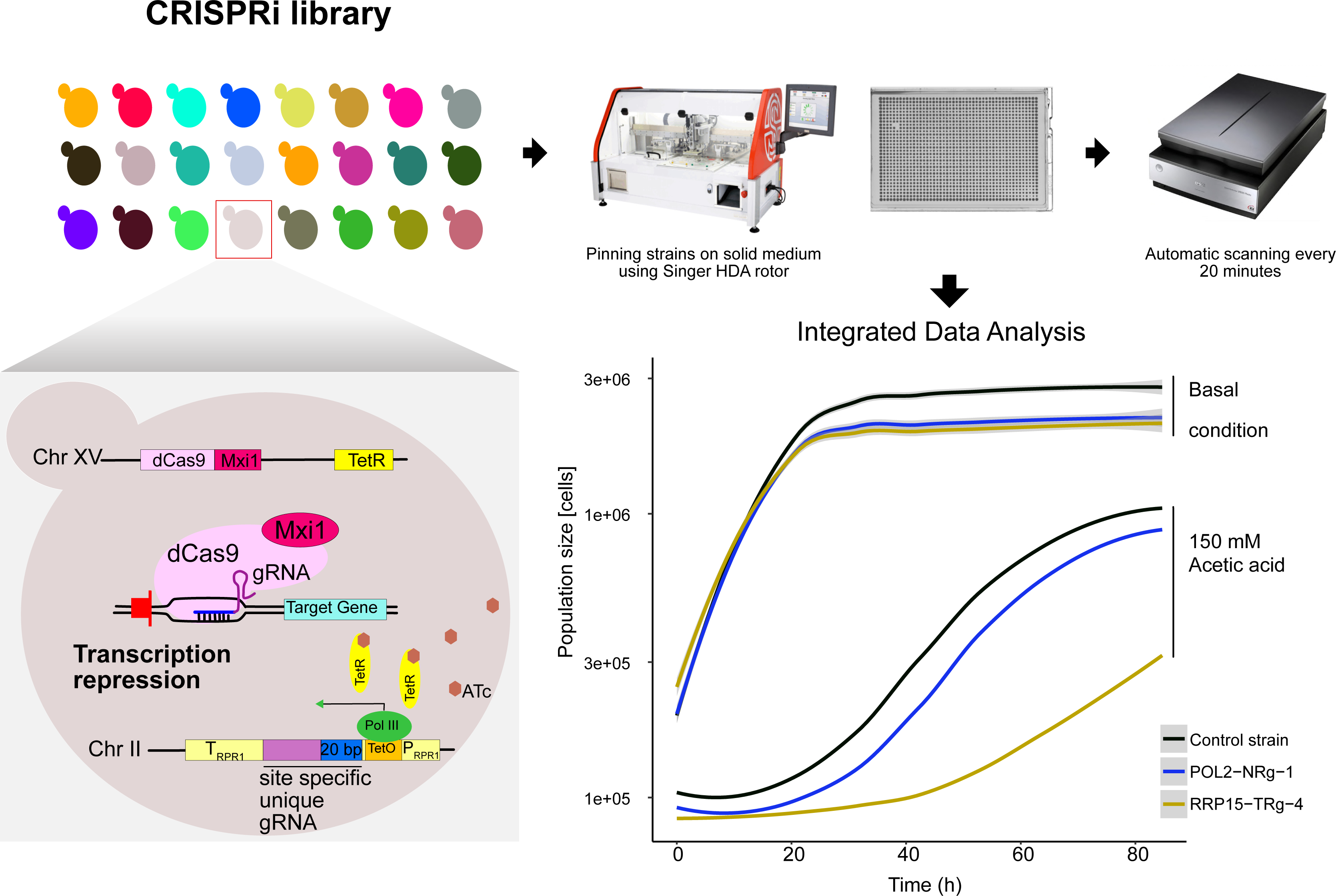
A constitutively expressed *dCas9-Mxi1* and the tetracycline-regulatable gRNA expression system induces transcription repression of essential or respiratory growth-essential genes. Each strain in the library was phenotyped individually for growth on solid medium with 150 mM acetic acid or in basal medium lacking acetic acid, using the Scan-o-matic platform.

**Fig. 2.**
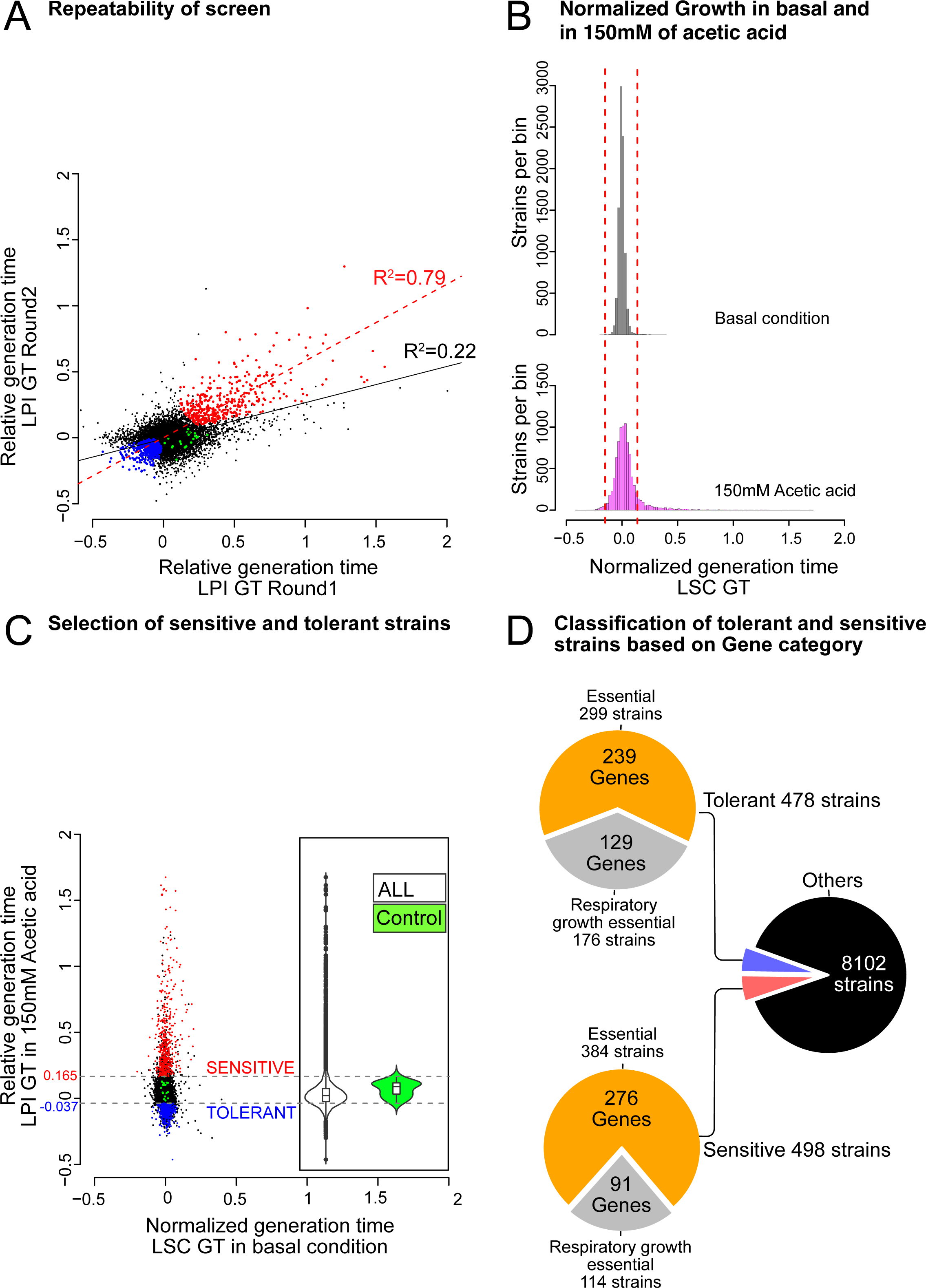
The CRISPRi strains showed minor phenotypic variation in basal condition and large phenotypic variation under acetic acid stress. A: Scatterplot displaying the reproducibility of the two Scan-o-matic screenings. The mean of the three LPI GT replicates of each strain is plotted, control strains in green, acetic acid sensitive strains in red, acetic acid tolerant strains in blue and remaining strains in black. The linear regression for the data of all strains is displayed with a black line and for the acetic acid sensitive and tolerant strains with a red line. B: Histogram of the normalized generation time of each CRISPRi strain in basal condition (grey) and at 150 mM of acetic acid (magenta). Strains outside the two red dashed lines, have generation times that are 10% shorter or 10% longer than the control strain. C: Scatterplot showing the normalized generation time of each CRISPRi strains in basal condition and relative generation time in medium with 150 mM acetic acid. Each point indicates the mean of all the replicates (n=6, when some of the replicates failed to grow, n=3-6). The data of the CRISPRi control strains is indicated green, of acetic acid sensitive in red, of acetic acid tolerant in blue and all other strains in black. The LPI GT threshold is indicated with a gray dashed line. Inset: the violin-plots display the spread and the distribution of the LPI GT data for all CRISPRi strains (ALL), and LPI GT values of CRISPRi control strains (Control). D: Overview of number of strains and genes identified as acetic acid tolerant or sensitive.

The CRISPRi strains showed limited phenotypic effects in basal condition and the generation time of 8,958 strains (99% of the strains of the library) was within ± 10% of the generation time of the control strain (Fig. 2B). Only 92 strains (1%) displayed complete growth inhibition in basal condition.

### CRISPRi-based gene repression imposed large phenotypic effects under acetic acid stress

In contrast to basal medium, large variations in generation time were observed among the CRISPRi strains at 150 mM of acetic acid (Fig. 2B and C). A great proportion of the CRISPRi strains displayed slower growth in response to acetic acid, with 1,040 strains (≈11%) having >10% higher generation time than the control strain. It was also clear from the growth curves that strains in acetic acid medium exhibited a rather long lag-phase before growth was resumed (Fig. 1). Still, 133 strains (≈1%) displayed a >10% shorter generation time than the control strain in response to acetic acid (Fig. 2B). In conclusion, the addition of acetic acid to the growth medium had a great impact on the growth of many of the strains in the CRISPRi library. The raw data and all the subsequent analytical output for all strains in the library are available in the supplementary .xlsx file, in Table S1 and S2.

**Table 1:** CRISPRi targeting of genes related to vesicle, organelle or vesicle transport induced acetic acid sensitivity.

| Gene | Gene description | Nr. of gRNAs | Change in generation time |
| --- | --- | --- | --- |
| Genes encoding COPI and COPII vesicle coating |  |  |  |
| *SEC27* | beta' subunit of COPI vesicle coat | 5 | +19 to +140% |
| *SEC21* | gamma subunit of COPI vesicle coat | 3 | +57% to complete inhibition |
| *RET3* | zeta subunit of COPI vesicle coat | 3 | +21 to +30% |
| *SAR1* | Regulation of COPII vesicle coating formation | 3 | +36 to +191% |
| *SEC23* | Regulation of COPII vesicle coating formation | 3 | +31 to +50% |
| *SEC24* | COPII vesicle cargo loading | 2 | +27 or +72% |
| *SEC13* | Facilitation of COPII vesicle budding | 2 | +25 or +119% |
| *SEC12* | Regulation of COPII vesicle coating formation | 1 | +19% |
| *SEC31* | Facilitation of COPII vesicle budding | 1 | +35% |
| *YPT1* | Facilitation of COPII vesicle budding | 1 | +39% |

| Genes encoding SNARE proteins |  |  |  |
| --- | --- | --- | --- |
| *YKT6* | v-SNARE protein | 2 | +22 or +76% |
| *BET1* | v-SNARE protein | 1 | Complete inhibition |
| *BOS1* | v-SNARE protein | 1 | +18% |
| *TLG1* | t-SNARE protein | 2 | +17% or +45% |
| *SED5* | t-SNARE protein | 2 | +30% or +197% |
| *SEC17* | involved in SNARE complex disassembly | 1 | +29% |
| *SEC22* | R-SNARE protein, assembles into SNARE complex with Bet1p, Bos1p and Sed5p | 1 | +15% |
| Vacuolar membrane ATPase complex proteins / GTPases required for vacuolar sorting |  |  |  |
| *VPS45\** | Essential for vacuolar protein sorting and also involved in positive regulation of SNARE complex assembly | 3 | +32% to +206% |
| *VMA3* | Proteolipid subunit c of the V0 domain of vacuolar H(+)-ATPase | 2 | +19% to +41% |
| *VMA7* | Subunit F of the V1 peripheral membrane domain of V-ATPase | 1 | +50% |
| *VMA11* | Vacuolar ATPase V0 domain subunit c | 1 | +18% |
| *VPS1* | GTPase required for vacuolar sorting | 2 | +71% to Complete inhibition |
| *VPS4* | AAA-ATPase involved in multivesicular body (MVB) protein sorting | 1 | +21% |
| *VPS36* | Involved in ubiquitin-dependent sorting of proteins into the endosome | 1 | +50% |
| *VPS53* | Required for vacuolar protein sorting | 1 | +48% |

### Integrative data-analysis connected yeast essential genes to acetic acid tolerance and sensitivity

In order to study gene-specific effects on acetic acid tolerance/sensitivity, we constructed relative generation times (LPI: log phenotypic index) where growth in acetic acid was put in relation to growth in basal medium. Thus, those strains that exhibited a general growth-defect and grew poorly in both media were not identified as specifically sensitive to acetic acid. Eleven % of all the strains (i.e. 954 strains, including 108 strains that did not grow in acetic acid) in the library had an increased relative generation time, while 19% of all the strains (1,704) had a decreased relative generation under acetic acid stress (Fig. 2C). A combined statistical (false discovery rate adjusted P-value ≤ 0.1) and effect size threshold was applied, which allowed the identification of 959 strains (corresponding to 665 genes) as acetic acid sensitive or tolerant (Fig. 2D). Out of these, 478 strains with gRNAs targeting a total of 370 genes had a significantly decreased relative generation time (Fig. 2D and Table S3), and thereby displayed acetic acid tolerance. The decrease in relative generation time seen was relatively small with only few strains showing a higher level of improvement, with RPN9-TRg-4 (targeting *RPN9*, encoding a regulatory subunit of the 26S proteasome; 27% improvement) and RGL1-NRg-7 (targeting *RGL1*, encoding a regulator of Rho1p signaling; 18% improvement) being the most acetic acid tolerant strains identified. A total of 498 strains, with gRNAs targeting a total of 367 genes, displayed acetic acid sensitivity (Fig. 2D and Table S4). Out of these, 17 strains that grew well in basal condition were completely inhibited (or the strains grew extremely slowly, generation time > 48h) in the presence of 150 mM acetic acid. The range of sensitivities was rather wide and the relative generation time for 34 strains was greater than 2-fold compared to the control strain, with ARC40-NRg-3 (targeting *ARC40*, encoding a subunit of the ARP2/3 complex; 219% extension) and VPS45-NRg-4 (targeting *VPS45*, encoding a protein essential for vacuolar protein sorting; 206% extension) being the most acetic acid sensitive strains. Thus, a rather large number of CRISPRi strains showed an altered response to acetic acid, where about half showed increased sensitivity and half increased tolerance.

### Growth in liquid media and qPCR expression analysis validated the large-scale phenomics results

To validate the phenomics data obtained from cultures grown on solid medium, the growth of 183 strains (including sensitive and tolerant strains as well as some controls), was analyzed also in liquid medium. In the liquid validation experiment both 150 mM and 125 mM acetic acid media were included, as the phenotypic effects were seen to be more drastic in liquid compared to solid medium. A high proportion of the strains did not grow at all in liquid medium at 150 mM, the concentration that was used in the screen on solid medium. The relative generation time in liquid medium showed a strong correlation (*r* = 0.86) with the corresponding Scan-o-matic data for growth on solid medium (Fig. 3, for representative growth curves of selected strains in liquid medium, see Fig. S1). Linear regression showed that 73% (R^2^ = 0.73, F-test P-value < 2.2e-16) of the phenotypic variation between these two independent experimental methods can be explained by the linear model. It should be noted that some strains can display, for biological reasons, different growth responses on solid and liquid media (20). We concluded that the data from the large-scale screen on solid medium was in excellent agreement with the liquid growth analysis.

**Fig. 3.**
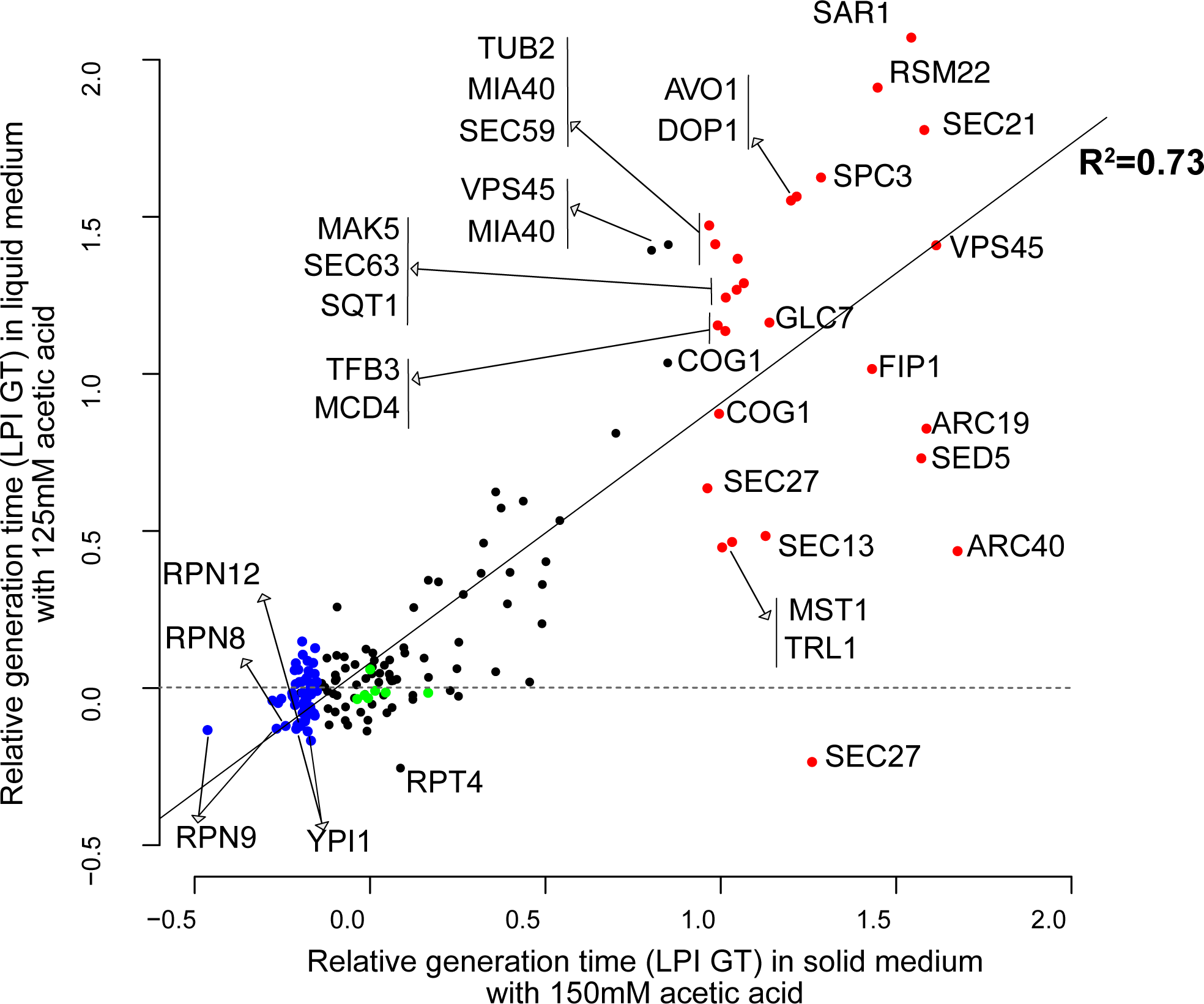
Scatterplot of the relative performance of the strains in liquid medium with 125 mM of acetic acid and in solid medium with 150 mM acetic acid (Scan-o-matic screening). The linear regression of the data is displayed with a black line. The mean of the three LPI GT replicates of each strain is plotted, control strains in green, acetic acid sensitive strains in red, acetic acid tolerant strains in blue and remaining strains in black. The names of the genes repressed in the tolerant or sensitive strains are indicated in the plot.

The initial screen on solid medium selected tolerant and sensitive strains only based on changes in growth rate (generation time). In addition to determination of generation time, the growth analysis in liquid media also allowed detailed analysis of growth lag and biomass yield. A sharp reduction in biomass yield was observed with increasing acetic acid stress (Fig. S2). During growth in liquid media, the generation time and yield of the strains showed a strong negative correlation both at 125 (r = - 0.91) and 150 mM (r = - 0.84) acetic acid, thus slow growth correlated with low yield during the cultivation. On the other hand, neither generation time nor yield correlated with the lag phase indicating that the length of the lag-phase is an independent physiological feature under acetic acid stress. The lag phase of strains grown in the presence of acetic acid was much longer compared to growth in basal medium, whereas the changes in generation time determined were less pronounced between the two types of media. An overview of the relative performance of the strains characterized in liquid medium is demonstrated using a heatmap in Fig. S3.

To investigate the relationship between the level of transcriptional repression of the target genes and the observed phenotypes, qPCR was performed for a selected set of strains with different generation times. The chosen strains had gRNAs targeting *RPN9*, *RPT4, GLC7* or *YPI1* (Fig. 4, S4). For most strains, different levels of repression of the target gene was observed using different gRNAs. For strains with gRNAs targeting *RPN9* or *GLC7*, the phenotype observed (faster growth in the case of *RPN9* and slower growth for *GLC7*) showed strong correlations with the reduction of expression levels of the target genes (*r* = 0.94 and *r* = -0.79 for *RPN9* and *GLC7*, respectively). The expression of *GLC7* in strains with the gRNAsGLC7-TRg-2 and GLC7-NRg-4 was strongly down-regulated (by ≈93% and ≈82%), and these two strains were also the most sensitive to acetic acid (+133% and +39% in relative generation time, Fig. 4C). For strains with gRNAs targeting *RPT4* or *YPI1*, there was no clear correlation between the change in expression levels and generation times (Fig. 4B and D).

**Fig. 4.**
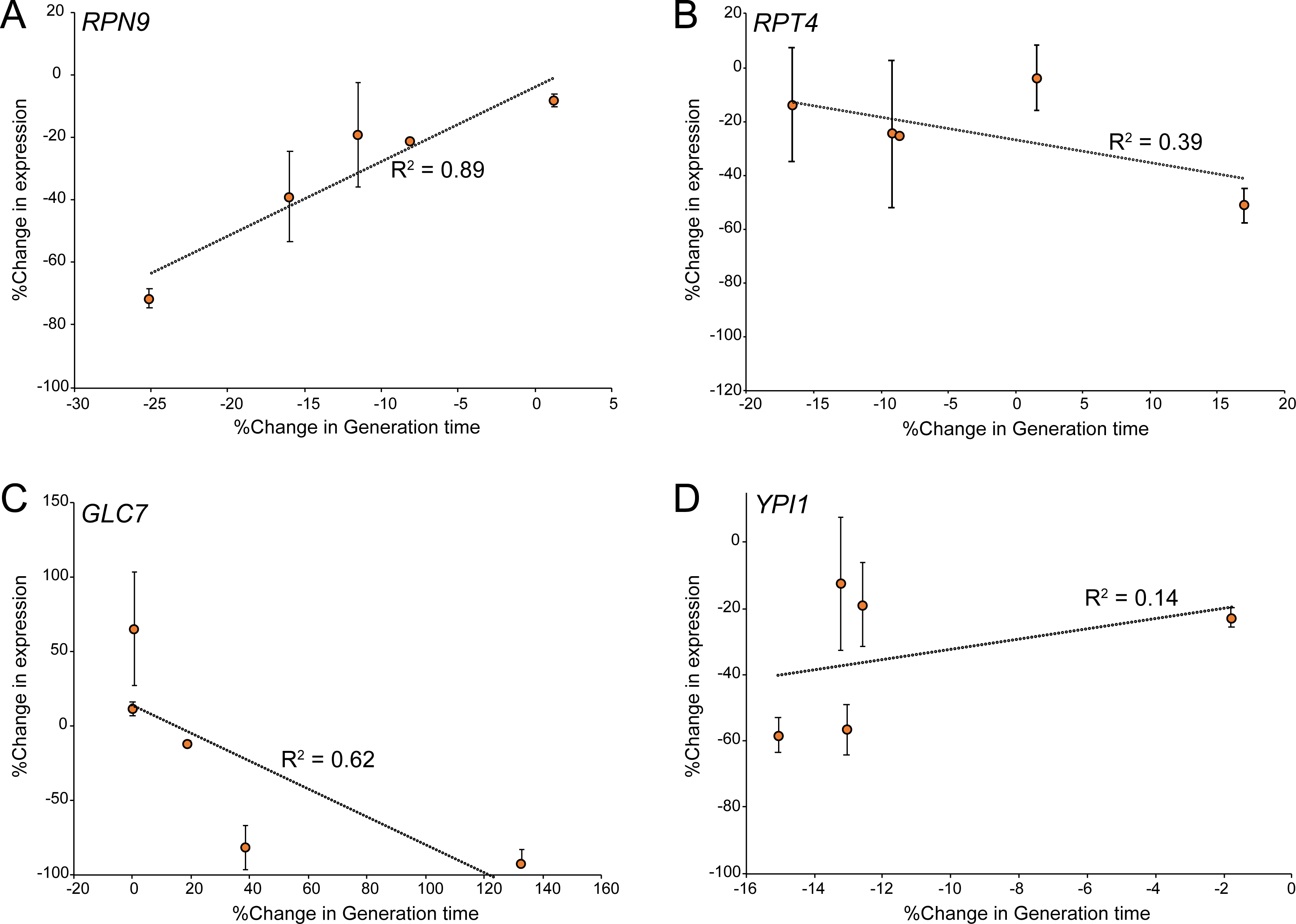
Percentage change in expression compared to the control strain of target genes at 125 mM of acetic acid in liquid medium in relation to percentage change in relative growth of selected CRISPRi strains compared to the control strain in solid medium with 150 mM of acetic acid. The gRNA of the strains targeted *RPN9* (A), *RPT4* (B), *GLC7* (C) or *YPI1* (D). The individual points on the plot represent different gRNAs targeting the same gene. The expression of the target gene was normalized against the geometric mean of the reference genes *ACT1* and *IPP1*. See Fig. S4 for qPCR data.

### Membrane bound organelles and vesicle mediated secretory pathways are of particular importance under acetic acid stress

Individual repression of 367 genes in 498 strains resulted in acetic acid sensitivity. Out of those genes, 276 are generally essential (represented by 384 strains) and 91 are respiratory growth essential genes (represented by 114 strains) (Fig. 2D, Table S4).

Gene Ontology (GO) enrichment analysis of genes for which repression imposed acetic acid sensitivity, indicated that a fully functional bounding membrane of different organelles is of great importance to handle acetic acid stress in *S. cerevisiae* (adjusted P-value = 0.00033, Fig. 5). The Golgi apparatus, endoplasmic reticulum (ER), vesicular structures such as the endosome, the vacuole and the organelle-associated intracellular transport pathways were found to be of particular importance (Fig. S5 and S6). Furthermore, several genes involved in vesicle mediated transport were enriched (adjusted P-value = 5.40E-05). Many strains with gRNAs targeting genes encoding the vacuolar membrane ATPase or GTPases required for vacuolar sorting (*VMA3*, *VMA7*, *VMA11*, *VPS1, VPS4*, *VPS36*, *VPS45* or *VPS53*) were found to be sensitive to acetic acid (Table 1). Moreover, the transport of luminal and membrane protein cargoes between the ER-Golgi segment of secretory pathway using COPI and COPII coated vesicles appeared crucial for growth under acetic acid stress. Strains with gRNAs targeting genes encoding beta’ (*SEC27*), gamma (*SEC21*) and Zeta (*RET3*) subunits of the COPI vesicle coat displayed severe sensitivity to acetic acid (Table 1). Similarly, CRISPRi repression of several genes that encode components involved in the regulation of COPII vesicle coating formation (*SEC12, SAR1, SEC23*), COPII vesicle cargo loading (*SEC24*), and components that facilitate COPII vesicle budding (*SEC31, YPT1, SEC13*) showed significant acetic acid sensitivity (Table 1).

**Fig. 5.**
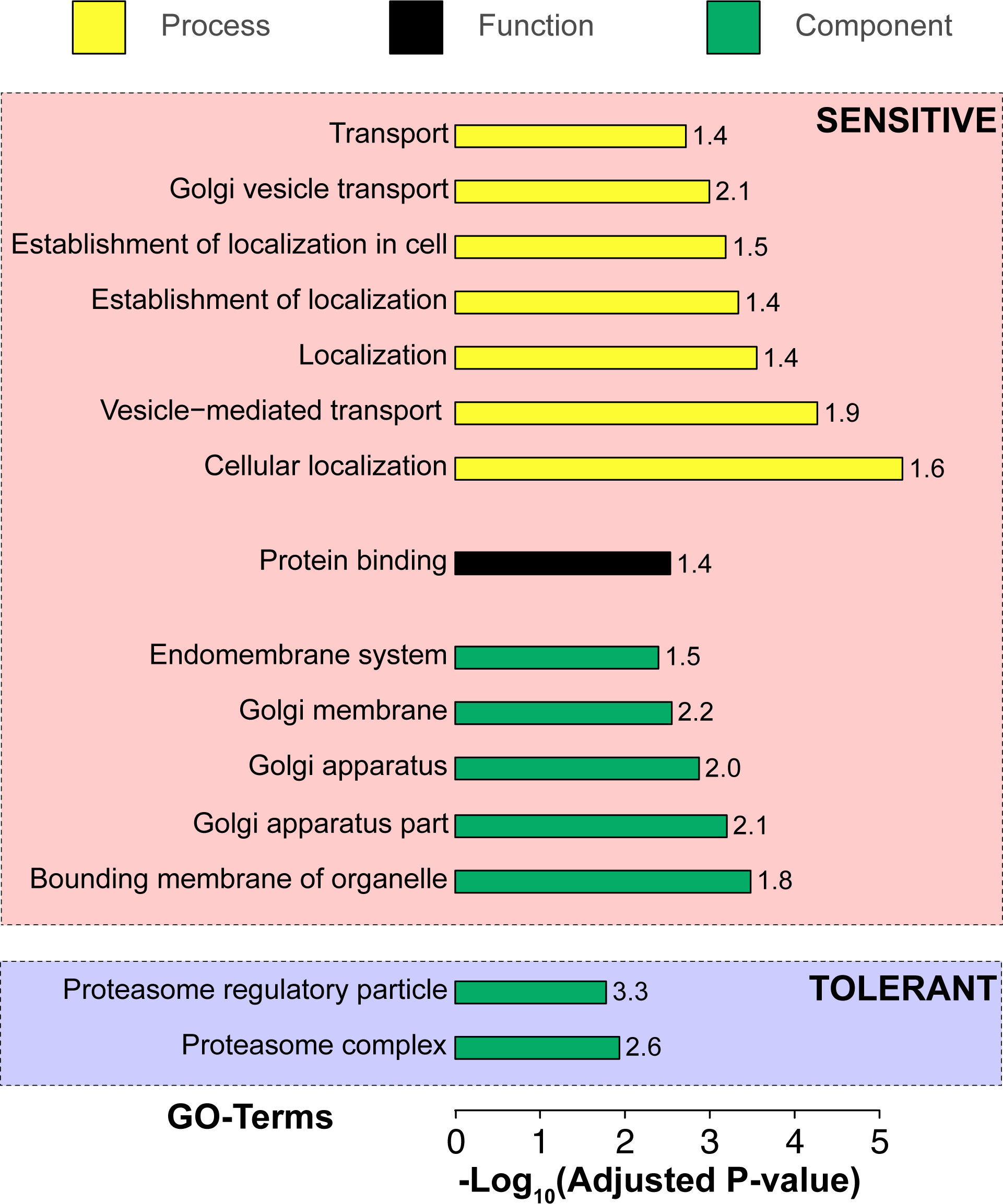
Functional and gene ontology enrichment analysis of genes repressed in acetic acid sensitive and tolerant CRISPRi strains. GO terms connected to biological process, genetic functions and cell components are indicated using yellow, black, and green colored bars, respectively. The negative log10-transformed Bonferroni-corrected P-value (Kruskal–Wallis test) is plotted on the X-axis. Enrichment factors (ratio of the observed frequency to the frequency expected by chance) for each GO term are displayed on the top of each bar.

In addition to COPI and COPII vesicle coating, our results also elucidated the importance of SNARE proteins, which mediate exocytosis and vesicle fusion with membrane-bound compartments. Our study included strains with gRNAs targeting 14 out of 24 known genes encoding SNARE proteins in *S. cerevisiae*. CRISPRi repression of 8 out of those 14 genes induced significant acetic acid sensitivity. In particular, CRISPRi repression of genes encoding v-SNARE proteins (proteins that are on the vesicle membrane) or t-SNARE proteins (proteins that are on the target membrane that the vesicles are fused to) increased the relative generation time in the presence of acetic acid (Table 1). We conclude that organelles and vesicle transport were highly enriched among sensitive strains, much in line with earlier reported features that are important for normal growth in acetic acid (21, 22).

### Repression of *YPI1*, involved in the regulation of the type I protein phosphatase Glc7, induced acetic acid tolerance

Accumulation of the storage carbohydrate glycogen has earlier been reported to be critical for growth under acetic acid stress (23, 24). *GLC7* encodes a type 1 protein phosphatase that contributes to the dephosphorylation and hence activation of glycogen synthases (25). We found that 3 out of 5 strains with gRNAs targeting *GLC7* showed significant acetic acid sensitivity, increasing the relative generation time by 16-120% (Fig. 4C). On the contrary, 5 strains with gRNAs targeting *YPI1*, a gene which has been reported to be involved in the regulation of Glc7, displayed significant acetic acid tolerance and reduced the relative generation time by 6-14% (Fig. 4D). The data obtained from solid medium was supported by data of strains growing in liquid medium, where one strain with a gRNA targeting *GLC7* was included. This strain showed significant acetic acid sensitivity (219% increment of relative generation time and 42% longer lag phase) at 125mM. In contrast, 3 out of 4 *YPI1* strains that were included in liquid growth experiment showed significant acetic acid tolerance (11-13% reduction in relative generation time and 3-11% reduction in lag phase in liquid medium with 125mM of acetic acid). In summary, our data give support for that Ypi1 acts as a negative regulator of Glc7 under acetic acid stress, and that it plays an important role during growth in acetic acid conditions, possibly by affecting the accumulation of glycogen.

### The proteasome regulatory subunits have a major role in acetic acid tolerance

Two GO-terms, i.e. “proteasome complex” and “proteasome regulatory particle”, were significantly enriched in the GO-analysis of the 370 genes that when repressed by the CRISPRi system displayed increased acetic acid tolerance (Fig. 5). Most of the genes connected to these GO terms encode subunits of the 19S regulatory particles (RPs) of the 26S proteasome (Fig. 6, S7). Among these were 6 genes (i.e. *RPN3, RPN5, RPN6, RPN8, RPN9, RPN12*; Table 2) encoding subunits for the RP lid assembly. The CRISPRi targeting of *RPN9* was most prominent with 5 out of 8 gRNAs inducing a significant decrease in the relative generation time, and multiple gRNAs targeting *RPN6* and *RPN5* also induced acetic acid tolerance (Table 2). Overall, the different gRNAs for these different RP lid assembly genes reduced the relative generation time in the range of 2 - 27% (Table 2).

**Fig. 6.**
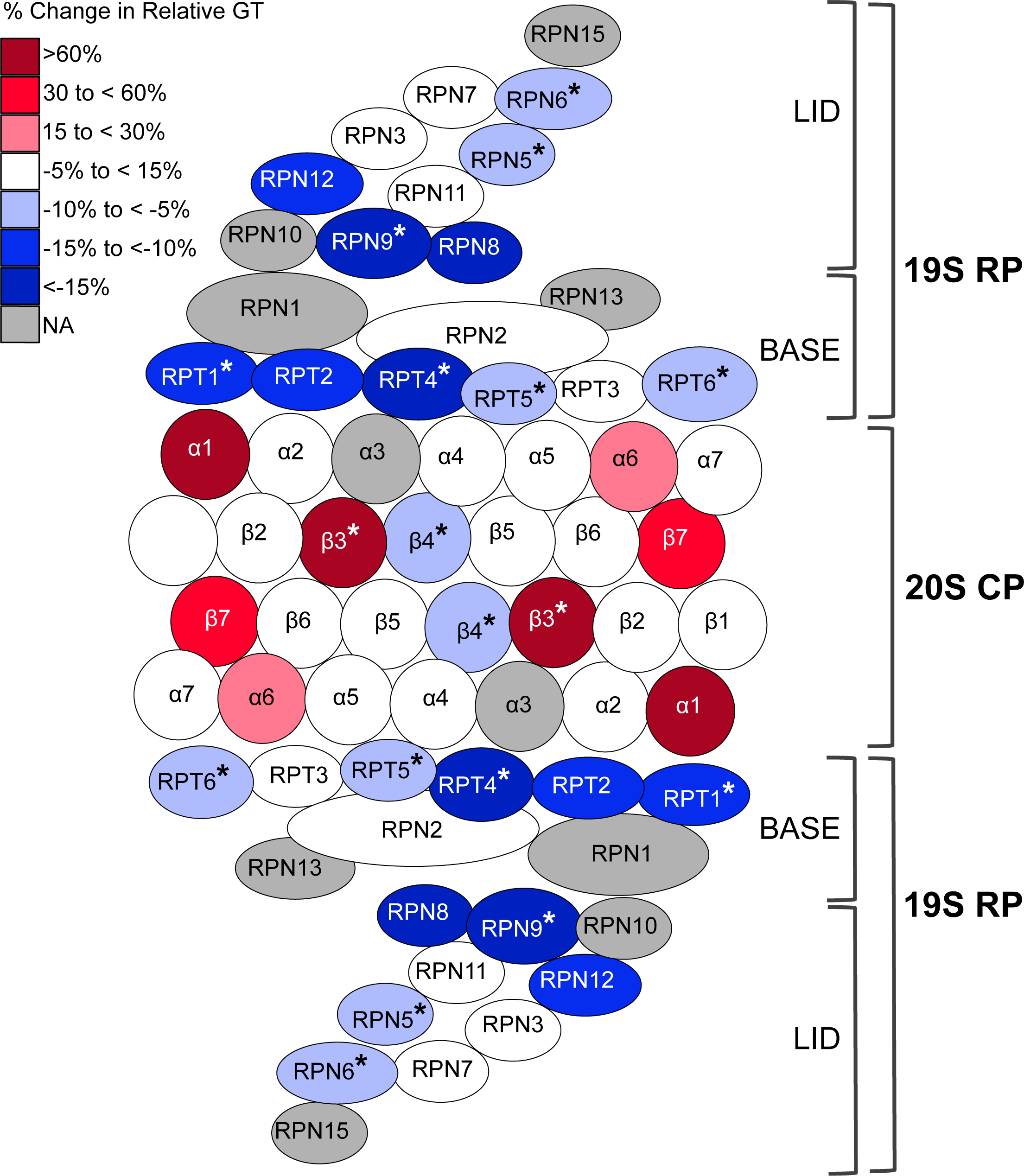
CRISPRi repression of genes encoding subunits of 26S proteasomal complex induced acetic acid tolerance (mainly genes encoding proteins of the 19S proteasomal regulatory particle lid and the base subcomplex, displayed with blue circles) or sensitivity (genes of the 20S core particle, displayed with red circles). The color in each subunit display only the most dominant phenotype (i.e. significant and highest in effect size) obtained by CRISPRi repression of the gene encoding that subunit. Subunits encoded by genes not included in the strain collection are displayed in grey and subunits for which CRISPRi repression with multiple gRNAs induced the dominant phenotype are indicated with an asterisk. The schematic representation of the relative positions of the subunits in the proteasome complex are inferred from the Cryo-EM structure reported by Luan et al. (66).

**Table 2:** CRISPRi targeting of genes encoding proteins of the 19S proteasomal regulatory particle lid and the base subcomplex induced acetic acid tolerance.

| Gene | Gene description | Nr. of gRNAs | Change in generation time |
| --- | --- | --- | --- |
| Proteasome 19S Regulatory particles LID complex |  |  |  |
| *RPN9* | Non-ATPase regulatory subunit of the 26S proteasome lid | 5 | -5% to -27% |
| *RPN6* | non-ATPase regulatory subunit of the 26S proteasome lid; required for the assembly and activity | 3 | -4% to -8% |
| *RPN5* | non-ATPase regulatory subunit of the 26S proteasome lid | 2 | -2% or -6% |
| *RPN3* | non-ATPase regulatory subunit of the 26S proteasome lid | 1 | -3% |
| *RPN8* | non-ATPase regulatory subunit of the 26S proteasome lid | 1 | -15% |
| *RPN12* | non-ATPase regulatory subunit of the 26S proteasome lid | 1 | -14% |
| Proteasome 19S Regulatory particles BASE complex |  |  |  |
| *RPT1* | ATPase of the 19S regulatory particle | 3 | -5% to -12% |
| *RPT4* | ATPase of the 19S regulatory particle | 2 | -11% to -16% |
| *RPT5* | ATPase of the 19S regulatory particle | 2 | -3% to -4% |
| *RPT6* | ATPase of the 19S regulatory particle | 2 | -6% to -7% |
| *RPT2* | ATPase of the 19S regulatory particle | 1 | -11% |
| Proteasome 20S Core particle |  |  |  |
| *SCL1* | Alpha 1 subunit of the 20S proteasome | 1 | +74% |
| *PRE5* | Alpha 6 subunit of the 20S proteasome | 1 | +15% |
| *PRE4* | Beta 7 subunit of the 20S proteasome | 1 | +31% |
| *PUP3* | Beta 3 subunit of the 20S proteasome | 2 | +30% to +63% |

The performance of ten strains with gRNAs targeting subunits of the 19S regulatory particle lid complex was also characterized in liquid media. Both strains with gRNAs inducing tolerance and gRNAs failing to give a measurable phenotype on solid medium were included. Most of the strains (4 out of 6) identified as tolerant on solid medium (with gRNAs targeting *RPN9* or *RPN12*) also showed significant acetic acid tolerance in liquid medium, with 8-12% reduction in relative generation time and 4-8% reduced lag phase at 125 mM of acetic acid (Fig. 7A).

**Fig. 7.**
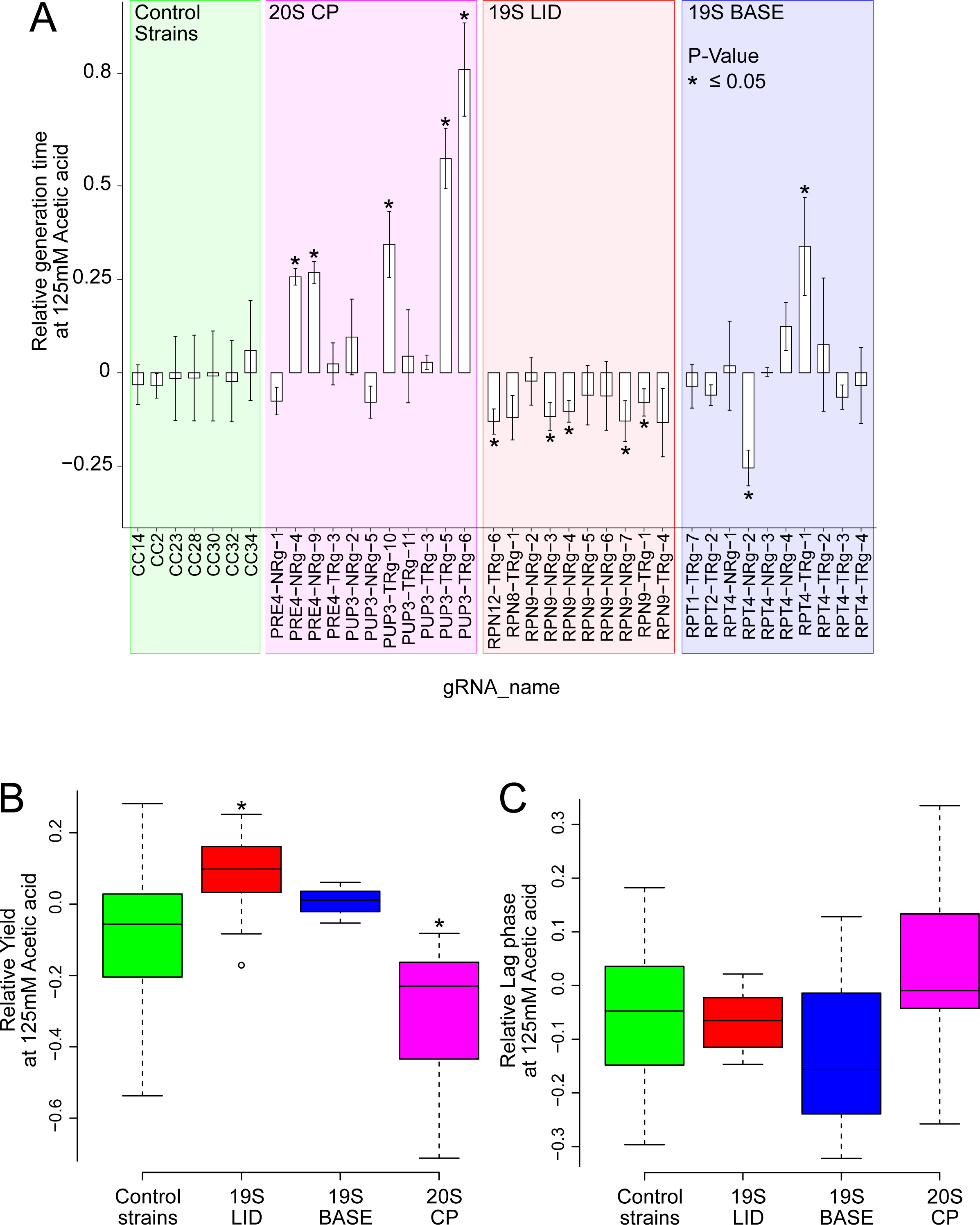
A: Relative growth in liquid medium of CRISPRi strains with gRNAs targeting genes encoding proteasomal subunits (20S CP; core particle, 19S lid or 19S base) and the control strains. The relative generation time of all strains (A) and biomass yield (B), and lag phase (C) of the acetic acid tolerant strains is shown.

In addition to acetic acid tolerance achieved by targeting the lid of the 19S regulatory particle, several CRISPRi strains targeting genes encoding subunits of the 19S RP base assembly showed significant acetic acid tolerance (Fig. 6 and Table 2). A reduction of 3-12% of the relative generation time was observed for strains with gRNAs targeting the RP base assembly subunits *RPT1*, *RPT2*, *RPT4*, *RPT5* or *RPT6*. The fitness benefit of targeting *RPT4* was confirmed in liquid medium, where the strain RPT4-NRg2 (Fig. 7A) had a 22% reduced relative generation time at 125 mM of acetic acid.

In contrast to the increased tolerance seen when targeting the 19S regulatory particle, CRISPRi targeting of genes encoding the 20S proteasome predominantly led to acetic acid sensitivity (Fig. 6). The relative generation times were increased by 15-74% in strains with gRNAs targeting *SCL1*, *PRE5*, *PRE4* or *PUP3* (Table 2). This trend was confirmed in liquid medium, where 6 out of 11 strains with gRNAs targeting genes encoding 20S proteasomal subunits showed significant acetic acid sensitivity (Fig. 7A). Thus, our data indicates that the proteasome and its different sub-parts play critical and differential roles in regulating growth in medium with acetic acid.

## DISCUSSION

### Bioengineering of essential genes in yeast using the CRISPRi technology

A number of large-scale, systematic gene-by-phenotype analyses of essential genes have previously been performed, by phenotyping either heterozygous deletion mutants or strains carrying temperature sensitive alleles (26-29). Nonetheless, the use of heterozygous deletion mutants is limited by haplosuffiency, as one copy of a gene often is adequate for the normal function of diploids (30). Moreover, temperature-dependent side-effects may influence the results when studying thermosensitive alleles (28, 31).

In previous studies where the CRISPRi technology was applied for massive genotype– phenotype mapping in *S. cerevisiae* (9, 11-13), the strains were pooled, and screened for competitive growth. Although competitive growth assays have the advantage of throughput, they come with a major weakness; the nutrient-specific advantage for cells/strains with shorter lag phase is amplified. Single cell analysis has showed massive heterogeneity in lag-phase within clonal populations of *S. cerevisiae* (32), which may introduce noise in the outcome of competitive growth assays. Moreover, the characterization of a population enriched after a specific time provides merely an endpoint observation. In the previously described competitive growth assays of whole genome CRISPRi libraries (11, 13) the genes identified to give beneficial phenotypes when repressed, have not been essential. This is likely due to the phenotypes of strains with altered expression of essential genes not being as pronounced as the phenotypes of the strains becoming enriched or due to the alteration in expression being detrimental. Often the genetic or environmental effects on cellular fitness are relatively small (33, 34), and thus highly accurate measurement methodologies are required to capture subtle differences in growth phenotypes. Therefore, we used the phenomics platform Scan-o-matic (20), to individually grow each of the >9,000 strains of a CRISPRi strain library. The generation time of each strain was generated from high-resolution growth curves without the influence/competition from other strains.

During growth in basal condition, we found that most of the CRISPRi strains grew with a generation time similar or just slightly slower compared to the generation time of the control strains. In medium with acetic acid, there was a great variability between the strains, some growing faster and as expected, many growing much slower. Only about 1% of the strains of the library did not grow in basal condition. This in line with what Smith et al. (12) observed when growing the pooled strains in YPD medium; after 10 doublings the DNA barcodes associated with 170 strains dropped below background. Our qPCR profiling of selected genes of strains during mid-exponential growth showed that both at basal condition and under acetic acid stress, different levels of repression was achieved by targeting the same gene with different gRNAs (Fig. 4). For the tested genes, we observed that the repression of expression was more pronounced in basal medium compared to medium supplemented with acetic acid (Fig. S4), indicating that the repression by the CRISPRi system may be influenced by the environmental condition. High concentrations of acetic acid are known to cause an increased lag phase (35). We observed that several of the strains scored for a change in growth rate also displayed defects or improvements on the length of the lag-phase, while some did not (Fig. S3).

### CRISPRi targeting vesicle, organelle or vesicle transport encoding genes causes acetic acid sensitivity

Previous large-scale screens of strains have identified many genes with widely diverse functions, the deletions of which increased the susceptibility of yeast to acetic acid (22, 36). In line with our findings, Sousa et al. (22) reported that deleting genes involved in vesicular traffic from the Golgi to the endosome and the vacuole increased sensitivity to acetic acid. In addition, endocytic inhibition has been observed in response to acetic acid and other environmental stressors (37). Many of the acetic acid sensitive strains in our study had gRNAs targeting genes encoding different proteins involved in the formation and activity of COPI and COPII vesicles or SNARE proteins (Table 1). The COPI and COPII vesicles transport proteins between the ER and the Golgi (reviewed by Szul and Sztul (38)), whereas SNARE proteins mediate exocytosis and vesicle fusion with different membrane-bound compartments (reviewed by Han et al. (39)). It has been reported that acetic acid causes ER stress and induces the unfolded protein response, as misfolded proteins accumulate in the ER (40). An earlier study, screening the deletion strain collection reported ER, Golgi, and vacuolar transport processes as important for resistance to a vast collection of small molecules or environmental stress conditions, including acetic acid treatment (41).

The deletions of genes encoding the vacuolar membrane ATPase complex (*VMA2-8*, *13*, *16*, *21*, *22*) has been shown to decrease the tolerance to acetic acid (22, 36), presumably as cells struggle to maintain a neutral cytosolic pH (42). Similarly, single gene deletions of VPS genes (encoding a GTPases required for vacuolar sorting) have been shown to result in a drastically enhanced sensitivity to acetic acid and a drop in intracellular pH (43). In line with these studies, we found strains with gRNAs targeting several vacuolar ATPase related genes (encoding VMA and VPS complexes; Table 1) to be among the sensitive strains, highlighting the importance of the vacuole in response to acetic acid stress.

### Regulation of genes involved in glycogen accumulation influence acetic acid tolerance

Glycogen serves as a fuel reserve for cells and accumulates when growth conditions deteriorate as a means of adapting to stress such as nutrient-, carbon- or energy-limitation (44), or acetic acid treatment (23, 24). Glycogen is produced from glucose-6 phosphate via glycogen synthases that are activated by dephosphorylation by e.g. the Glc7 phosphatase (25). Hueso et al. (45) demonstrated that overexpression of a functional, 3’-truncated version of the *GLC7* gene improved acetic acid tolerance. In our study, 3 strains with gRNAs targeting *GLC7* showed strong acetic acid sensitivity (Fig. 4C). Ypi1 was initially reported to be an inhibitor of Glc7 (46), while it was later shown to positively regulate Glc7 activity in the nucleus (47). Overexpression of *YPI1* has been shown to reduce glycogen levels (46). Our study showed that downregulation of *YPI1*, encoding a regulatory subunit of the type I protein phosphatase Glc7, conferred acetic acid tolerance. Five strains with gRNAs targeting *YPI1* displayed a significant decrease in generation time when subjected to acetic acid, and the downregulation of *YPI1* in these CRISPRi strains was confirmed by qPCR (Fig. 4D).

In light with the fact that both Ypi1 and Glc7 have many different roles in maintaining cell homeostasis beyond glycogen synthesis, we propose that a CRISPRi-mediated repression of *YPI1* may be favorable for the cells under acetic acid stress, likely due to increased glycogen levels in the cells. Similarly, we suggest that CRISPRi strains where *GLC7* is repressed may have decreased intracellular glycogen content, thus rendering them more sensitive to acetic acid. Still it may be that other regulatory roles of Ypi1 and Glc7 are behind the acetic acid resistance/sensitivity identified for some of the CRISPRi strains and determination of this needs further study.

### Adapting proteasomal degradation of oxidized proteins to save ATP increases acetic acid tolerance

While the best-known function of the proteasome is ATP-dependent protein degradation through the 26S ubiquitin-proteasome system, the unbound, ATP-independent, 20S proteasome is the main protease responsible for degrading oxidized proteins (reviewed by Reynes et al. (48)). The 26S proteasomal complex consists of one 20S core particle and two 19S regulatory particles that are further divided into lid- and base-assemblies. In our study, many of the strains with increased acetic acid tolerance had gRNAs targeting genes encoding subunits of the 19S regulatory particle of the proteasome (Fig. 6 and 7A).

Many studies report on accumulation of reactive oxygen species (ROS) under acetic acid stress and reactive oxygen species are well known to cause protein oxidation and even induce programmed-cell-death in cells upon acetic acid stress (reviewed by Guaragnella et al. (49)). Yeast under oxidative stress respond to the accumulation of ROS with a decrease in cellular ATP concentration (50). Acetic acid that enters the cell dissociates to protons and acetate ions at the near-neutral cytosolic pH and the charged acetate ions are unable to diffuse through the plasma membrane and thus accumulate intracellularly (reviewed by Palma et al. (42)). Therefore, acetic acid stress, in particular pumping out excess protons from the cytosol to the extracellular space by H^+^-ATPase pumps in the plasma membrane and from the cytosol to the vacuole by the vacuolar H^+^-ATPases, causes a reduction in ATP (42). Moreover, the accumulation of ROS has been reported to induce a metabolic shift from glycolysis towards the pentose phosphate pathway in order to increase the production of NADPH, an essential cofactor to run the antioxidant systems, which leads to reduction in ATP generation (51). Consequently, ATP conservation by reducing the activity of ATP-dependent processes could offer yeast a fitness benefit against acetic acid stress.

The 20S core particle on its own performs ubiquitin- and ATP-independent degradation of proteins. Under acetic acid stress, ROS accumulation triggers protein oxidation that leads to protein unfolding (52). The inner proteolytic chamber of the 20S core particle is only accessible to unfolded proteins and moderately oxidized proteins are ideal substrates for the 20S proteasome (53-55). We hypothesize that the repression of subunits of the 19S regulatory particle increases the abundance of free 20S core particles, which offers the cell an alternative to the ATP expensive 26S proteasome mediated protein degradation. In line with this, it has been reported that even mild oxidative stress reversibly inactivates both the ubiquitin activating/conjugating system and the 26S proteasome activity but does not impact the functionality of the 20S core particle (56, 57). Therefore, an increased abundance of the 20S core particle alone in the strains where the CRISPRi system targets genes encoding subunits of the 19S regulatory particle could allow more efficient ATP-independent degradation of oxidized proteins, thus conferring yeast a fitness benefit during acetic acid stress (Fig. 8).

**Fig. 8.**
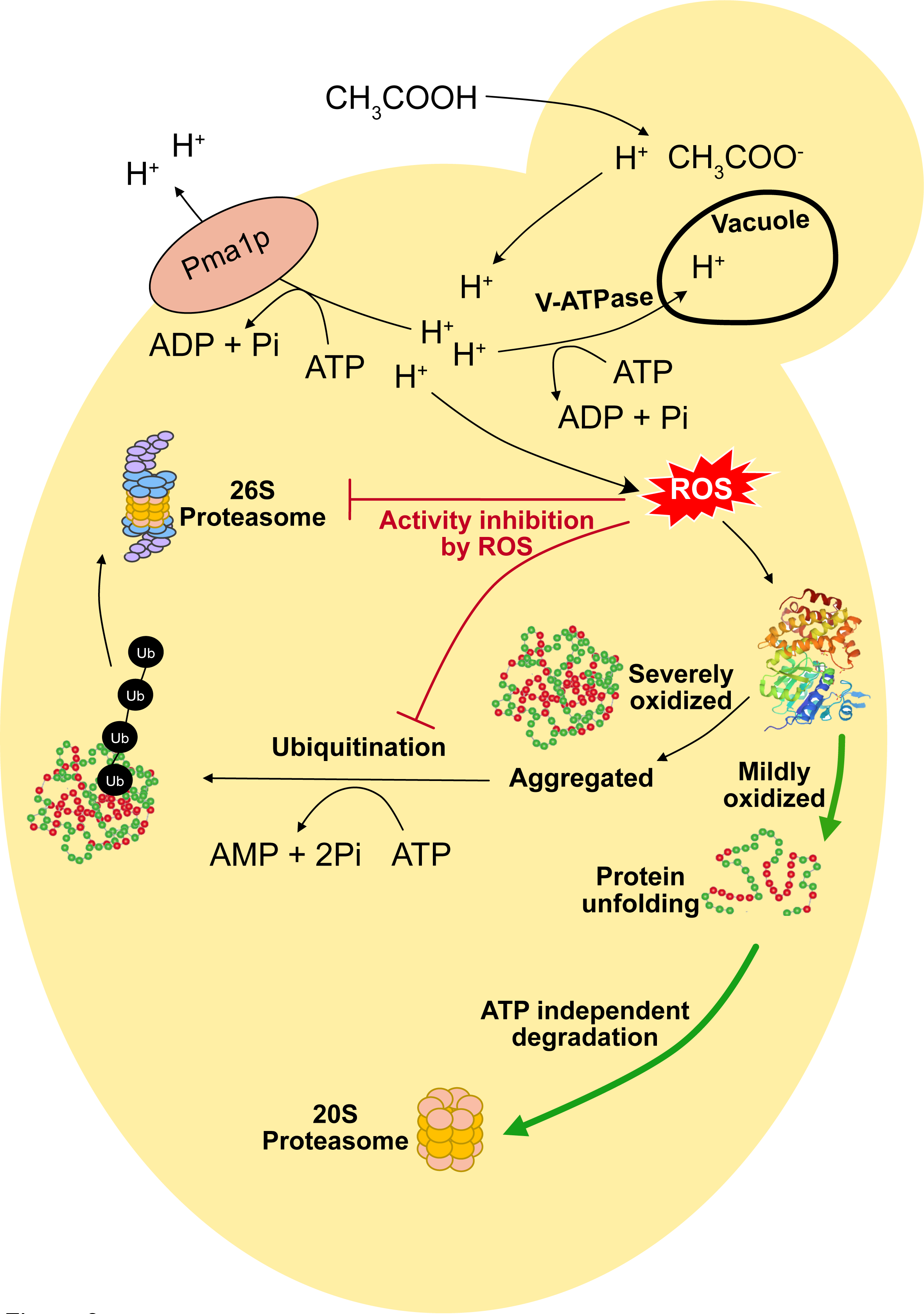
Overview of the response of the cells towards acetic acid stress based on CRISPRi targeting of essential genes. The cells are starved of ATP due to ATP expensive processes such as the elevated action of H^+^-ATPase and V-ATPase pumps. Therefore, we hypothesize that the downregulation of subunits of 19S RP increases the abundance of 20S CP, which offers the cell an alternative to the ATP expensive 26S proteasome mediated protein degradation. This in turn gives yeast a fitness benefit under oxidative stress induced by acetic acid. ROS = reactive oxygen

A total of five CRISPRi strains with gRNAs targeting *RPN9* (encoding a subunit of the 19S regulatory lid-assembly) had significantly decreased generation times in medium supplemented with acetic acid (Table 2). This gives confidence that downregulation of *RPN9* provides a means to improve acetic acid tolerance. Previously, an *rpn9* mutant with defective assembly of the 26S proteasome and reduced 26S proteasome activity, was shown to be more resistant to hydrogen peroxide that is a common stressor used to enforce oxidative stress (58). Moreover, this *rpn9* mutant was able to degrade carbonylated (oxidized) proteins more efficiently than the wild type strain and it displayed an increased 20S-dependent proteasome activity (58). In our study, we observed that the yields of strains with gRNAs targeting the 19S lid or base of the proteasome was increased for strains growing in acetic acid whereas the yield of strains with gRNAs targeting the 20S CP of the proteasome was decreased (Fig. 7B). It seems plausible that the repression of genes encoding subunits of the 19S lid lead to decreased ATP-expensive 26S activity and that this ATP saving contributed to a concomitant increment in biomass.

Our qPCR results showed that the level of repression of *RPN9* or *RPT4* (encoding a subunit of the 19S regulatory base-assembly) was greatly dependent on the gRNA of the strains (Fig. 4A and B). For *RPN9* there was a strong correlation between expression level and acetic acid tolerance, indicating that fine tuning the 20S and 26S proteasomal regulation could be an efficient strategy to bioengineer acetic acid tolerant industrial yeast strains (Fig. 4A). In line with this, a recent study showed that the downregulation of *RPT5* (encoding a subunit of the 19S base-assembly) induced tolerance against oxidative stress (59). In our study, down-regulation of *RPT4* was for 3 out of 5 strains with gRNAs targeting this gene shown to improve acetic acid tolerance (Fig. 4B and S3B). Nonetheless, the generation time of RPT4-TRg-1 with a clear repression of *RPT4* was increased. We argue that a too strong repression of an essential gene is likely to be detrimental, highlighting the need for a fine-tuned expression when engineering tolerance. While off-target effects of gRNAs as well as gRNAs failing to give a phenotype is a known challenge of the CRISPRi technology, screening several strains with different gRNAs and identifying multiple strains with similar phenotypes gives confidence in a phenotype being a result of the gene repression itself (12). In our study, a total of 28 strains with gRNAs targeting proteasomal genes were identified as tolerant or sensitive (Table 2), which gives great confidence for us to elaborate on the role of the proteasome during acetic acid stress.

In conclusion, our study identified many essential and respiratory growth essential genes that regulate tolerance to acetic acid. CRISPRi-mediated repression of genes involved in vesicle formation or organelle transport processes led to severe growth inhibition during acetic acid stress, emphasizing the importance of these intracellular membrane structures in maintaining cell vitality. The data also suggests that an increased activity of the ATP-independent protein degradation by the 20S core is an efficient way of counteracting acetic acid stress. This mechanism may ensure ATP savings, allowing proton extrusion and an increased biomass yield. A fine-tuned expression of proteasomal genes could be a strategy for increasing stress tolerance of yeast, leading to improved strains for production of biobased chemicals.

## MATERIALS AND METHODS

### Yeast strain library

The CRISPRi strain library (12) used in this study contains 9,078 strains, each of which has an integrated dCas9-Mxi1 repressor (14). The strains also contain a tetracycline-regulatable repressor (TetR), where the TetR controls a modified Pol III promoter (TetO-PRPR1) that drives the expression of unique gRNAs (Fig. 1). Thus, the gRNAs are expressed in the presence of the inducing agent, anhydrotetracycline (ATc). Each strain in this library expresses a unique gRNA that in combination with dCas9-Mxi1, targets 1,108 of the 1,117 (99.2%) essential genes (30) and 505 of 514 (98.2%) respiratory growth essential genes (60, 61) in *S. cerevisiae* (Fig. S8A and B). For most of the genes (1,474 out of 1,617), there are at-least three and up to 17 strains (mean ≈ 5), with different gRNAs targeting the same gene in the library (Fig. S8C). 93% of the unique gRNAs were designed within 200 bp upstream of the transcription starting site of the respective target gene (Fig. S8D). Depending on the targeting location of the gRNA in the promoter, genetic repression ranging from very strong to weak can be achieved (8). This produces strains that under ATc induction have different levels of repression of the same gene relative to the native expression level. Moreover, 20 strains in the CRISPRi library have gRNAs that are non-homologous to the *S. cerevisiae* genome and function as control strains (Fig. S8B). The CRISPRi strains were stored in YP Glycerol stock solution (17% Glycerol (v/v), 10 g/l Yeast extract, 20 g/l Bacto-peptone). The whole collection was kept in 24 microtiter plates (MTP 384 well format). Unless otherwise mentioned, all chemicals were purchased from Merck.

### ATc titration in YNB medium

Synthetic defined medium was used to identify acetic acid-specific effects, excluding compounds present in rich medium that might confound the interpretation of our data. To obtain appropriate gene suppression in our set-up, we adjusted the concentration of ATc in relation to what had been proposed earlier for rich-media liquid cultures (12). The concentration of ATc sufficient to induce high level of gRNA expression in the CRISPRi strains growing on YNB agar media, was determined by a qualitative spot test assay with selected strains (Fig. S9A). These strains were selected based on the competitive growth assay of the CRISPRi library in liquid YPD medium with and without 250 ng/ml of ATc by Smith et al. (12). This study showed that growth of the strains with gRNA targeting the essential genes *ACT1* (ACT1-NRg-5: TTAAACAAGAGAGATTGGGA, ACT1-NRg-8: ATTTCAAAAAGGAGAGAGAG), *VPS1* (VPS1-TRg-1: GCCGGGTCACCCAAAGACTT) and *SEC21* (SEC21-NRg-5: GTCGTAGTGAATGACACAAG) was nearly or completely inhibited, as these essential genes, targeted by the gRNAs of the strains, were strongly repressed. These strains, as well as two control strains i.e. Ctrl_CC11 (CC11: CCCAGTAGCTGTCGGTAGCG) and Ctrl_CC23 (CC23: AGGGGTGCTAGAGGTTTGCG) were grown on synthetic defined Yeast Nitrogen Base (YNB) agar medium, (YNB; 1.7 g/l Yeast Nitrogen Base without amino acids and ammonium sulfate (BD Difco), 5 g/l ammonium sulphate, 0.79 g/l Complete Supplement Mixture with all amino acids and nucleotides (Formedium), 20 g/l glucose, 20 g/l agar, succinate buffer i.e. succinic acid 10 g/l and sodium hydroxide 6 g/l), in the presence of 0, 2.5, 5, 7.5, 10, 12.5 or 25 μg ATc /ml. A stock solution (25 mg/ml in dimethyl sulfoxide; DMSO) was used to achieve the different ATc concentrations. The final concentration of DMSO in the media was adjusted to 0.1% (v/v). The pre-cultures for the spot assay were grown in liquid YNB medium for 48 h, after which 3 µl drops from serial dilutions (10^-1^, 10^-2^, 10^-3^, and 10^-4^) of a cell suspension of OD600 1 were spotted on solid YNB medium with different concentrations of ATc and incubated at 30°C for 48 h. We found that 2.5 μg/ml of the gRNA inducer ATc was sufficient to elicit growth defects on solid medium for strains with gRNAs targeting the essential genes *ACT1*, *VPS1* or *SEC21*. The growth of these strains was incrementally inhibited up to near complete inhibition at 7.5 μg /ml ATc (Fig. S9A). In contrast, the growth of the control strains (strains expressing gRNAs with no genomic target) remained unimpeded even at 25 μg/ml ATc (Fig. S9A) and we therefore used 7.5 μg /ml ATc in our screen of the CRISPRi library.

A liquid ATc titration assay was done in 200 μl liquid YNB medium with 0, 0.25, 1, 2, 3, 5, 7.5, 10, 15, or 25 μg ATc /ml in a Bioscreen C MBR device (Fig. S9B). The strains were pre-cultured in YNB medium for 48 h. A separate pre-culture was used to inoculate each replicate at a starting OD600 of approximately 0.1. In order to avoid uneven oxygen distribution, the plastic cover of the bioscreen plate was replaced with a sterile sealing membrane permeable to oxygen, carbon dioxide and water vapor (Breathe-Easy®, Sigma-Aldrich). Strains were grown in continuous shaking for 75 h during which automated spectrophotometric readings were taken every 20 min. The raw data was calibrated to actual OD600 values and smoothed before the growth lag, generation time, and growth yield were estimated using the PRECOG software (62). All growth experiments were performed at 30°C.

### Media preparation for high-throughput phenomics

Solid YPD (10 g/l yeast extract, 20 g/l bacto-peptone, 20 g/l glucose, 20 g/l agar) medium was used to re-grow the CRISPRi collection from the -80°C storage, and also to grow the pre-cultures. Growth phenotypes of all the CRISPRi strains in the library were evaluated in the basal condition i.e. solid YNB medium and in solid YNB supplemented with 150 mM acetic acid. ATc (7.5 μg/ml, as determined by the qualitative spot assay) was added to both media to induce gRNA expression. The acetic acid concentration used was determined by growing a sub-set of the CRISPRi strains (739 strains), that were pinned to the actual experimental format of 1,536 colonies per plate, on solid medium with different acetic acid concentrations (50, 75, 100, 150 mM). The largest phenotypic difference in growth between the strains was observed at 150 mM of acetic acid (Fig. S10), and this concentration was selected to be used in our screen. The final concentration of DMSO in the growth media was 0.03% (v/v) and the pH was adjusted to 4.5.

### High-throughput phenomics using Scan-o-matic

The high-throughput growth experiments were performed using the Scan-o-matic (20) phenomics facility at the University of Gothenburg, Sweden. The procedure is described here in short. A robot Singer ROTOR HDA was used for all replica pinning. First, the frozen -80°C stock of the CRISPRi library in 24 microtiter plates was pinned in 384-array format on solid YPD medium and then incubated at 30°C for 72 h in scanners imaging the plates. For each of the 24 plates, one pre-culture plate was prepared in 1,536-array format. For this purpose, 384 strains were pinned thrice so that each has 3 adjacent replicates. In this way 384x3 i.e. 1,152 positions in a 1,536 array was filled. All fourth positions i.e. the rest of the 384 positions were filled with a spatial control strain to normalize any spatial growth bias (Fig. S11). The Scan-o-matic system uses a dedicated algorithm that can normalize any spatial growth bias in the extracted phenotypes of the other strains using the growth data of this spatial control strain (20). Here, the control strain Ctrl_CC23 was used as the spatial control strain. The preculture plates were incubated at 30°C for 48 h before being used for the replica pinning on the experimental plates, that were placed in the scanners in a predefined orientation and incubated at 30°C. The plates were imaged automatically every 20 min for 96 h. Subsequently, image analysis by Scan-o-matic was performed and a growth curve was generated for each colony. Finally, absolute, and spatially normalized generation times were extracted for all replicates of each strains. The whole experimental process was repeated twice to generate 6 experimental replicate measurements for each strain in both the medium with 150 mM acetic acid and in the basal medium lacking acetic acid.

### Data analysis

R version 4.0.2 was used to perform all mathematical and statistical analysis. The analytical steps employed to identify essential or respiratory growth-related genes that lead to acetic acid sensitivity or tolerance when repressed are described below. Here we also explain terminologies used.

#### Normalized generation time (LSC GT) and batch correction

The normalized generation time obtained after the spatial bias correction gives the population doubling time of a strain colony relative to the spatial control strain on a log2 scale (20). This is referred to as the log strain coefficient for the generation time (LSC GT). In previous studies it was found that only few of the gRNAs targeting a specific gene can induce strong repression that results in a strong phenotypic effect (8, 12) and therefore most of the strains will display a phenotype similar to that of a control strains. Since we used the control strain Ctrl_CC23 as the spatial control strain, it was expected that the median LSC GT of all strains in an experimental plate would be close to zero. However, some variability in the dataset was still present due to unavoidable micro-environmental factors between plates and this caused a slight deviation of the median value of the LSC GT for some experimental plates. To correct for this batch effect, a plate-wise correction was conducted by subtracting the median of LSC GT values of all the individual colonies on a plate from the individual LSC GT values of the colonies growing on that plate. i.e. if strainX is growing on plate Z, the corrected LSC GT value for strainX was the following:

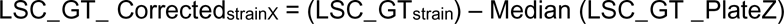

#### Relative generation time in the presence of acetic acid (LPI GT)

The growth of each CRISPRi strain was evaluated in two different conditions i.e. medium with 150 mM acetic acid (AA_150 mM_) and basal medium lacking acetic acid (Basal.condition). The relative performance of a strain in the presence of acetic acid compared to the basal condition was determined by subtracting the LSC_GT_ Basal.condition from the LSC_GT_ AA_150 mM_. This relative estimation, which gives the acetic acid specific effect on the generation time (GT) of a strain, is defined as the log phenotypic index (LPI GT, (63)), i.e. for strainX the LPI GT was calculated as the following;

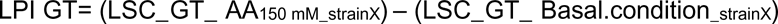

#### Statistical tests and P-value adjustment

Since it was expected that most strains would show only minor changes in generation time, here it is hypothesized that a phenotypic difference between a specific CRISPRi strain to the mean phenotypic performance of all the CRISPRi strains that falls within the interquartile range (IQR) of the complete dataset (i.e. having a LPI GT value between -0.024 to 0.075) would be zero, and any difference within the IQR to be just by chance. Therefore, formally our null hypothesis was the following:

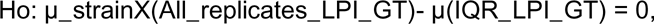

i.e. the difference between LPI GT mean of all replicates of strainX to the mean of the LPI GT dataset within IQR equals zero. The P-value for each strain in the library was estimated using Welch’s two sample two-sided t-test, which is an adaptation of Student’s t-test and produces fewer false positives (64). Moreover, this method remains robust for skewed distributions and large sample sizes. In this study, the mean LPI GT of 3392 strains displayed a significant (P-value ≤ 0.1) deviation from µ_[IQR_LPI_GT]_ when subjected to Welch’s two sample two-sided t-test (Fig. S12A). The P-values were corrected by the Benjamini-Hochberg method, also known as the false discovery rate (FDR) method (65). An adjusted-value threshold of ≤ 0.1 was set to select acetic acid tolerant or sensitive strains. Application of the FDR method (65) left 1258 strains below the adjusted P-value threshold of 0.1 (Fig. S12B). None of the control strains had an adjusted P-value below 0.1 (Fig. S12D).

An LPI GT threshold was applied for the selection of tolerant or sensitive strains. If a CRISPRi strain had an LPI GT_Mean that was greater than the maximum of the LPI GT_Mean of the control strains, then the strain was considered as acetic acid sensitive. Similarly, if a CRISPRi strain had an LPI GT_Mean less than the minimum of the LPI GT _Mean of the control strains, then the strain was considered as acetic acid tolerant. In this study, we observed that the range of the LPI GT_mean for control strain was between -0.037 and 0.166.

Therefore, acetic acid sensitive strain = µ_strain_ (LPI GT) > 0.166 and P.adjusted-value ≤ 0.1 acetic acid tolerant strain = µ_strain_ (LPI GT) < -0.037 and P.adjusted-value ≤ 0.1

Some CRISPRi strains that grew well in the basal condition but very poorly or not at all on the acetic acid experimental plates were identified. These strains were not subjected to any statistical analysis, but still added to the final list of acetic acid sensitive CRISPRi strains.

### Gene ontology (GO) analysis

GO term (process, function, and component) enrichment analysis of the gene lists of acetic acid tolerant and sensitive strains was performed against a background set of genes (all 1617 genes targeted in this CRISPRi library) using the GO term finder in the *Saccharomyces* genome database (Version 0.86) (https://www.yeastgenome.org/goTermFinder) and all GO term hits with p-value < 0.1 were identified.

### Data access

The R scripts used for analysis and the phenomics data generated in this project are available from https://github.com/mukherjeevaskar267/CRISPRi_Screening_AceticAcid. The raw image files of the Scan-o-matic projects can be requested for reanalysis from the authors.

### Growth of selected strains in liquid media

In order to validate the acetic acid sensitivity or tolerance observed for the CRISPRi strains in the Scan-o-matic screening, selected strains were grown in liquid YNB medium using the Bioscreen platform. The 48 most acetic acid sensitive and 50 most tolerant CRISPRi strains from the Scan-o-matic analysis were selected for the validation. Moreover, all CRISPRi strains with gRNAs targeting any of the following 12 genes: *RPT4, RPN9, PRE4, MRPL10, MRPL4, SEC27, MIA40, VPS45, PUP3, VMA3, SEC62, COG1*, were included making a total of 176 strains that were grown together with 7 control strains in liquid medium (raw data available in Table S5).

Briefly, the strains were pinned from the frozen stock into liquid YNB medium and grown at 30°C for 40 h at 220 rpm. This plate was used as the preculture and separate precultures were prepared for each independent culture. The strains were grown in liquid YNB medium (basal condition) and in liquid YNB medium supplemented with 125 or 150 mM of acetic acid. For each strain, 3 independent replicates were included for each growth condition. Two μg/ml ATc was added to the media to induce gRNA expression. The final concentration of DMSO in the growth media was 0.008% (v/v) and the pH was adjusted to 4.5. The experimental method and subsequent phenotype extraction were the same as for the ATc titration experiment, except that the strains were grown for 96 h. Similar to Scan-o-matic, all downstream analysis was performed using R version 4.0.2.

### Expression analysis by qPCR

#### Strains

Expression analysis by qPCR was performed to detect mRNA expression of *RPN9*, *RPT4*, *GLC7* and *YPI1*. For each target gene, 5 strains (i.e. each with a different gRNA) that showed different degree of acetic acid tolerance/sensitivity in Scan-o-matic screening were selected. Three control strains (CC2, CC23, CC32) were included to estimate the expression of the target genes in the absence of CRISPR interference.

#### RNA preparation and cDNA synthesis

Cells were grown to mid-exponential phase in liquid YNB (basal condition) or YNB medium supplemented with 125 mM acetic acid, in the Bioscreen platform and collected by centrifugation at 2000xg, at 4°C for 3 min. The cell pellet was immediately frozen in liquid nitrogen. For each CRISPRi strain, 2 independent replicates and for each control strain 3 independent replicates were included. For RNA preparation, the pellet was dissolved in 600 µl lysis buffer (PureLink™ RNA minikit, Invitrogen) after which the cell-suspension was transferred into tubes containing 0.5 mm glass beads. Cells were lysed by shaking for 40 s at 6 m/s in a MP Biomedical FastPrep and then collected by centrifugation in a microcentrifuge for 2 min at 4 °C, at full speed. 370 µl of 70% ethanol was added to the resulting supernatant and total RNA was prepared using the PureLink™ RNA minikit (Invitrogen). The obtained RNA was treated by DNase (TURBO DNA-free™ Kit, Invitrogen) and cDNA synthesis was performed on 900 ng DNased RNA using the iScript™ cDNA Synthesis Kit (Bio-Rad).

### Measurement of gene expression

qPCR was performed using 2.5 ng cDNA and the iTaq™ Universal SYBR® Green Supermix (Bio-Rad) for detection. The expression of the target genes was normalized against the geometric mean of the reference genes *ACT1* and *IPP1*. Primer efficiencies were between 96 and 102% as determined by using different amounts of cDNA. For primer sequences see Table S6. The qPCR protocol was as follows: an initial denaturation at 95°C for 3 min, denaturation at 95°C for 20s, annealing at 60 °C for 20 s and elongation at 72°C for 30 s. In total 40 PCR cycles were run. For statistical analysis, an F-test was performed to determine the variance between all the replicates of the control strains and the replicates of a CRISPRi strain. Depending on this result, a two sample two tailed t-test assuming equal or unequal variance was performed for each strain and for a particular condition, where the null hypothesis was:

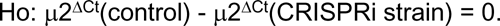

## ACKNOWLEDGMENTS

The Novo Nordisk Foundation (Grant number NF19OC0057685), The Royal Swedish Academy of Sciences and Chalmers Area of Advance Energy are acknowledged for financial support.

## AUTHOR CONTRIBUTIONS

Y.N. and A.B. conceptualized the project; Y.N, A.B. and V.M. designed the experimental and computational analysis; V.M. and U.L. performed the experiments; V.M. performed computational analysis; V.M., Y.N., A.B., and R.P.S. interpreted the results; V.M. and Y.N. wrote the initial draft paper; all authors revised the initial draft and wrote the final paper.

